# Molecular and Behavioral Effects of Infant Maltreatment across Generations in Rhesus Monkeys

**DOI:** 10.1101/791152

**Authors:** Torsten Klengel, Elyse L. Morin, Brittany R. Howell, Sara Bramlett, Dora Guzman, Jerrold S. Meyer, Kerry J. Ressler, M. Mar Sanchez

## Abstract

Childhood maltreatment is among the most robust risk factors for subsequent psychiatric and medical disorders, and data in humans and rodents suggest that effects of adverse childhood experiences may be transmitted across generations. Recent indications for biological processes underlying this transfer of experiential effects are intriguing; yet, their relevance in primates is inconclusive due to limitations of current studies. In this study, we bridge research in rodent models and humans with a natural non-human primate model on intergenerational effects of childhood maltreatment. Using a unique, well-controlled, randomized cross-fostering design in rhesus monkeys, we test the influence of ancestral maltreatment on molecular, neuroendocrine and behavioral outcomes in offspring, and show that childhood maltreatment results in transmission of information to the subsequent generation independent of behavioral transmission. We further demonstrate differences in the offspring longitudinal DNA methylation profile of the *FKBP5* gene, an important regulator of the hypothalamus-pituitary-adrenal axis in offspring of the maltreatment lineage compared to the control lineage. Finally, we show that differences in *FKBP5* methylation have functional effects on molecular, neuroendocrine and behavioral outcomes in offspring of the maltreatment ancestral line, even if the infants were never exposed to maltreatment nor interacted with their exposed ancestors. Although the molecular mechanism underlying our observations remains unknown, our data point to a potential germline-dependent effect. In summary, our data suggest that history of maltreatment in primates can induce molecular and behavioral changes in a subsequent generation independent of behavioral transmission that may influence risk for mental and physical diseases across generations.

## Introduction

Exposure to childhood maltreatment is one the most important risk factors for a variety of psychiatric disorders ^1,2^. However, both human studies and animal models suggest that effects of environmental experiences such as maltreatment extend beyond the affected individual, influencing phenotypes in subsequent generations. This sparked a controversial discussion on the prevalence of these phenomena, the underlying mechanisms and their biomedical relevance ^3–5^. Strong evidence exists for effects of environmental cues on offspring phenotypes when exposed during *in utero* development ^6–8^, or for the consequences of parental experience on offspring phenotypes mediated by parental behavior ^9^. More recently, experiments in rodents suggested that pre-conceptional exposure to chemicals ^10^, drugs ^11^, nutritional abnormalities ^12^ and stress ^13–15^ can prompt intergenerational effects or even lead to transgenerational inheritance of specific phenotypes ^16^. These events seem independent of *in utero* perturbations or behavioral transmission and point to possible germline-dependent mechanisms. Notably, some of the above studies have even demonstrated intergenerational transmission of phenotypes via *in vitro* fertilization. Although recent indications for similar effects with a focus on psychiatric phenotypes in humans are certainly intriguing, these studies are confounded by genetic, behavioral, nutritional, medical and socio-economic factors ^17–26^. In addition, multigenerational studies in humans are extremely lengthy and difficult to control. It thus remains unclear if such mechanisms exist in humans and to what extent they may influence physical and mental health.

Childhood maltreatment and other forms of early life stress (ELS) may cause alterations in long-term cellular programming of relevant pathways including stress response systems such as the hypothalamus-pituitary-adrenal (HPA) axis ^27–30^. Epigenetic processes, often broadly defined in the sense of cellular reprogramming, have been proposed as attractive mechanisms to explain the influence of environmental factors on genome function in general, and the regulation of gene expression in particular ^31,32^. However, there is an ongoing debate, particularly in primates, as to *if* and *how* environmental information can be transmitted or even inherited through the gametes across generations, potentially preparing the offspring generation for a similar environment through epigenetic adaptation ^16,33^. In this study, we focus on the epigenetic regulation of FK506 binding protein 5 (*FKBP5*), a co-chaperone of the glucocorticoid receptor (GR), and an important regulator of the HPA axis influencing the negative feedback control of cortisol and GR activation across species. Prior genetic and epigenetic studies repeatedly implicated *FKBP5* as a risk factor for psychiatric disorders in response to childhood trauma ^34–42^. In fact, we and others have previously shown that childhood maltreatment in humans lead to changes in DNA methylation within the *FKBP5* gene, a dysregulation of the negative feedback mechanism towards the GR, and subsequently a higher risk for stress- and trauma-related disorders ^42,43^.

Very similar to humans, non-human primate (NHP) infants exhibit drastic socioemotional and neural development over a protracted postnatal period during which the infant is critically dependent on parental care ^44–46^. Infant maltreatment occurs spontaneously in NHP species, both in captivity and in the wild, at rates similar to those seen in humans ^47–51^. In rhesus monkeys, infant maltreatment has been operationalized by the co-occurrence of physical abuse and rejection of the infant by the mother during the first 3 to 6 months of life, leading to infant pain and distress and elevations of stress hormones ^45,52–55^. Our group has extensively characterized the behavioral, neuroendocrine and neurodevelopmental impact of this naturalistic, translational NHP model of maltreatment. We previously demonstrated maltreatment effects on HPA axis hyperreactivity, telomere length, heightened anxiety and emotional reactivity, impulsive aggression and social deficits during development, effects which are consistent with chronic stress exposure ^46,55,56^. Further, infant maltreatment leads to increased amygdala volumes associated with higher emotional reactivity ^57^ and reduces the integrity of brain white matter tracts that connect cortico-limbic regions important for processing of sensory and emotional stimuli ^58^. Strikingly, infant maltreatment in NHPs is a relatively stable maternal trait, repeated with subsequent offspring and transmitted across generations ^48,59^. This matches similar observations in humans showing that parents with a history of experiencing maltreatment are more likely to show abusive behavior towards their own children compared to parents without this history ^60^. Previous studies revealed that cross fostering of infants from a maltreating line to a control line can often halt this vicious cycle, suggesting that the transmission of maternal maltreatment across generations is partially experiential and independent of genetic effects ^59^. However, prior studies using this model did not consider the possible non-genetic transmission of maltreatment-related phenotypes across generations and their relevance for disease risk. Importantly, models of ELS or early rearing experiences in NHP support the hypothesis that effects of nursery rearing, variable foraging demand stress, and variations in normal nurturing behavior, among others, may be transmitted across generations ^61–64^. Thus, we sought to use our ELS model with cross-fostering to examine the possibility of transmission of epigenetic risk in a well-controlled design. For this, we randomly assigned newborn macaques at birth from maltreating or competent care lineages to competent care mothers to specifically investigate the effect of pre-conceptional maltreatment in the maternal generation on the offspring epigenetic profile and related phenotypes. We carefully generated these animals with high genetic and social rank diversity, by choosing them from different social ranks, matrilines and paternities that were balanced across the experimental groups and sexes ^58,59,65^.

In summary, using a naturalistic model of ELS in NHPs, this study provides evidence for intergenerational, biological programming of *FKBP5* DNA methylation in response to ancestral maltreatment. Infant maltreatment in mothers, a pre-conceptional adverse experience, leads to a decrease in *FKBP5* DNA methylation, as well as HPA neuroendocrine and behavioral alterations in their offspring in the absence of direct interaction of the biological mother with the progeny (i.e. independent of maternal care). Our data highlight the long-term effects of childhood maltreatment across generations potentially increasing risk for psychiatric disorders.

## Results

### Study Sample

To investigate the contribution of ancestral maltreatment on the next generation molecular, neuroendocrine and behavioral phenotypes, we focus on infants born to maltreated (MALT) or control (CTRL) biological mothers that were randomly assigned and cross-fostered at birth to control foster mothers with a history of competent, nurturing maternal care (CTRL) following previous protocols ^58^, thus creating two separate groups: CTRL-to-CTRL and MALT-to-CTRL (**Figure 1**). A summary of descriptive data of the cohort divided by biological mother, foster mother and offspring is provided in **Table 1**. The statistical results are presented as mean ± SD throughout, unless otherwise indicated. Mean age (years) of the biological mothers was 9.93 ± 3.35 (MALT: 9.77 ± 3.86, CRTL: 10.06 ± 3.06, t(18)=0.18, p=0.85); mean age of the foster mothers was 10.43 ± 3.35 (MALT-to-CTRL: 9.69 ± 2.97, CRTL-to-CTRL: 11.04 ± 3.66, t(18)=0.89, p=0.38). Both, biological and foster mothers were experienced multiparas without significant differences with respect to parity (Biological MALT: 4.11 ± 2.76, Biological CTRL: 3.64 ± 2.5, t(18)=−0.40, p=0.69 and Foster MALT-to-CTRL: 3.78 ± 2.22, Foster CTRL-to-CTRL: 4.09 ± 2.81, t(18)=0.27, p=0.78) or gravida (Biological MALT: 4.33 ± 2.78, Biological CTRL: 4.18 ± 2.64, t(18)=0.13, p=0.90 and MALT to CTRL: 4.56 ± 2.56, CTRL to CTRL: 4.82 ± 3.13, t(18)=0.20, p=0.84). Importantly, biological mothers did not differ with respect to social rank distribution (MALT rank [high/middle/low]: 2/4/3, CTRL rank [high/middle/low]: 3/2/6 X^2^(2, N=20)=1.68, p=0.43). Similarly, foster mothers did not show differences in social rank (MALT to CTRL rank [high/middle/low]: 3/5/1, CTRL-to-CTRL rank [high/middle/low]: 2/3/6 X^2^(2, N=20)=4.11, p=0.13). Offspring groups did not differ in sex (MALT: M/F=3/6, CTRL: M/F=6/5, X^2^(1, N=20)=0.90, p=0.34) or rank movement between biological and foster mothers (MALT: [down/none/up]: 2/2/5, CTRL: [down/none/up]: 4/4/3, X^2^(2, N=20)=1.65, p=0.44). Thus, groups were balanced with respect to known relevant confounding variables.

**Figure 1.**
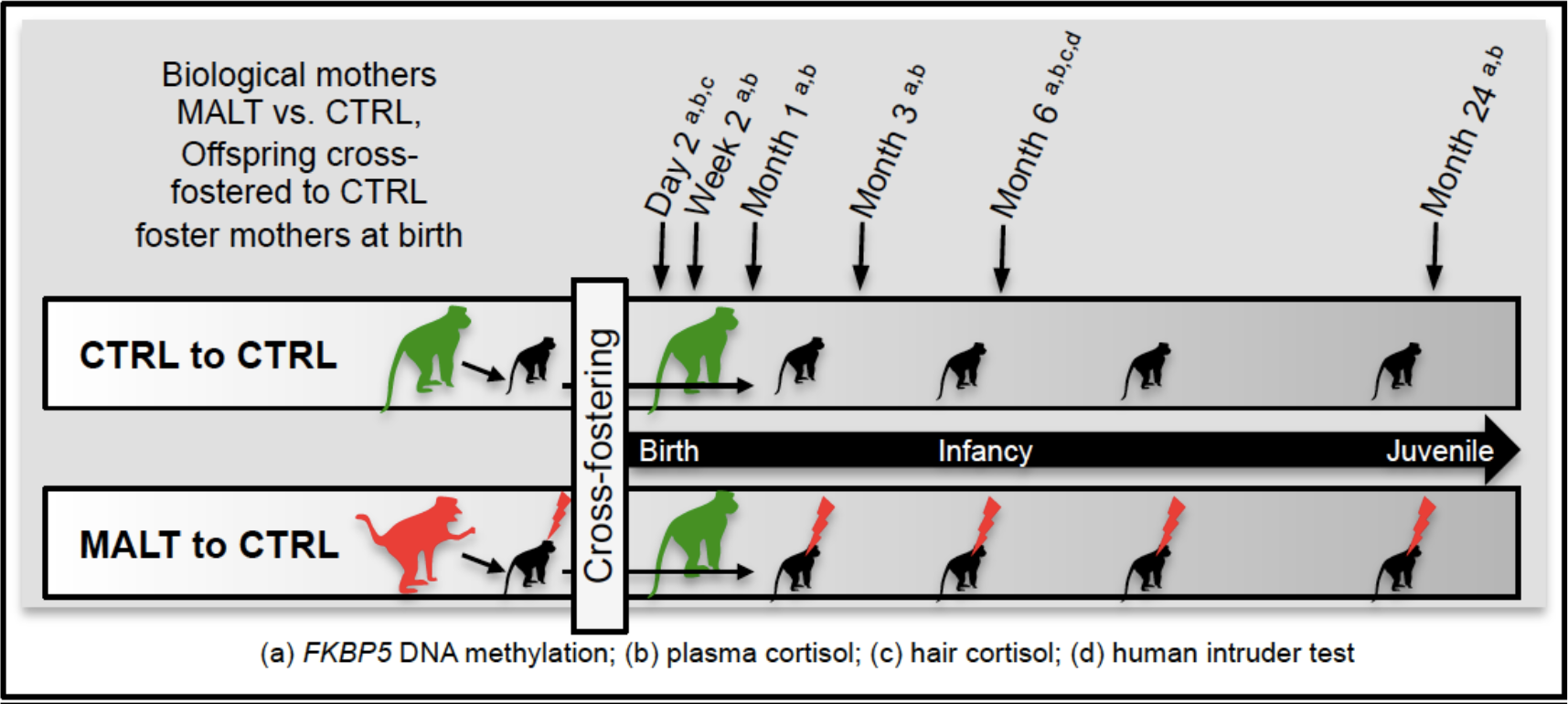
Experimental overview. In order to specifically investigate the effect of infant maltreatment in the ancestral generation, offspring from MALT and CTRL biological mothers were cross fostered after birth to CTRL foster mothers that provided competent care. We followed the offspring from birth to 24 months of life. EDTA blood was obtained at day 2, week 2, month 1, month 3, month 6, and month 24. Hair samples were obtained at day 2 and month 6. A human intruder test was performed at month 6.

**Table 1.** Demographic data of biological mothers, foster mothers and offspring. Demographic data for MALT and CTRL biological mothers, CTRL foster mothers, and offspring are shown. Groups were balanced with respect to important confounding variables.

| **Biological Mothers** | **MALT** | **Min** | **Max** | **SD** | **CTRL** | **Min** | **Max** | **SD** |  | **Combined** | **Min** | **Max** | **SD** |
| --- | --- | --- | --- | --- | --- | --- | --- | --- | --- | --- | --- | --- | --- |
| Total N | 9 |  |  |  | 11 |  |  |  |  | 20 |  |  |  |
| Mean Age (years) | 9.77 | 5.92 | 18.27 | 3.86 | 10.06 | 6.94 | 16.06 | 3.06 | t(18)=0.18, p=0.85 | 9.93 | 5.92 | 18.27 | 3.35 |
| Mean Gravida (N) | 4.33 | 1 | 9 | 2.78 | 4.18 | 2 | 11 | 2.64 | t(18)=-0.13, p=0.9 | 4.25 | 1 | 11 | 2.63 |
| Mean Parity (N) | 4.11 | 1 | 9 | 2.76 | 3.64 | 1 | 10 | 2.5 | t(18)=-0.40, p=0.69 | 3.85 | 1 | 10 | 2.56 |
| Social Rank (High/Middle/Low) | 2/4/3 |  |  |  | 3/2/6 |  |  |  | X<sup>2</sup>(2, N=20)=1.68 p=0.43 | 5/6/9 |  |  |  |
| Mean DNA methylation CpG5 (%) | 90.38<sup>(1)</sup> | 78 | 100 | 8.29 | 91.56<sup>(2)</sup> | 72 | 100 | 9.3 | t(15)=0.28, p=0.78 | 91 | 72 | 100 | 8.59 |
| **Foster Mothers** | **MALT** | **Min** | **Max** | **SD** | **CTRL** | **Min** | **Max** | **SD** |  | **Combined** | **Min** | **Max** | **SD** |
| Total N | 9 |  |  |  | 11 |  |  |  |  |  |  |  |  |
| Mean Age (years) | 9.69 | 6 | 14.02 | 2.97 | 11.04 | 6.94 | 17.82 | 3.66 | t(18)=0.89, p=0.38 | 10.43 | 6 | 17.82 | 3.35 |
| Mean Gravida (N) | 4.56 | 2 | 9 | 2.56 | 4.82 | 2 | 11 | 3.13 | t(18)=0.20, p=0.84 | 4.7 | 2 | 11 | 2.81 |
| Mean Parity (N) | 3.78 | 1 | 7 | 2.22 | 4.09 | 1 | 10 | 2.81 | t(18)=0.27, p=0.78 | 3.95 | 1 | 10 | 2.63 |
| Social Rank (High/Middle/Low) | 3/5/1 |  |  |  | 2/3/6 |  |  |  | X<sup>2</sup>(2, N=20)=4.11 p=0.13 | 5/8/7 |  |  |  |
| **Offspring** | **MALT** |  |  |  | **CTRL** |  |  |  |  |  |  |  |  |
| Total N | 9 |  |  |  | 11 |  |  |  |  | 20 |  |  |  |
| Sex (M/F) | 3/6 |  |  |  | 6/5 |  |  |  | X<sup>2</sup>(1, N=20)=0.9 p=0.34 | 9/11 |  |  |  |
| Social Rank Movement between Biological and Foster Mother (Down/None/Up) | 2/2/5 |  |  |  | 4/4/3 |  |  |  | X<sup>2</sup>(2, N=20)=1.65 p=0.44 |  |  |  |  |

Footnotes: <sup>(1)</sup> N=8 and <sup>(2)</sup> N=9

### Maternal phenotype and FKBP5 DNA methylation in the offspring

Similar to evidence shown in humans, maltreatment in monkeys spans generations with females exposed to maltreatment being at higher risk to maltreat their own offspring ^48,59,65^. To test the hypothesis that MALT in the biological mothers influenced cellular epigenetic programming in the offspring independent of maternal care, we measured *FKBP5* DNA methylation in intron 7, covering the genomic region investigated before in our human studies on childhood trauma ^42^ (**Supplemental Figure 1**). Focusing on the ancestral maternal phenotype (MALT vs. CTRL), linear mixed effect models show significant lower methylation in MALT offspring than CTRLs already present at birth at CpGs in and around the second glucocorticoid response element (GRE) of intron 7, which are orthologous to the sites found previously demethylated in the human *FKBP5* gene in childhood trauma studies ^26,35,42^. After correction for multiple testing, DNA demethylation at CpG5 remains significant (β=0.28, SE=0.08, p=0.003, p_Bonferroni_=0.018) (**Figure 2**). Thus, in light of the cross-fostering design, our data provide strong evidence for an intergenerational effect of maltreatment on offspring *FKBP5* methylation mediated though mechanisms independent of behavior, with potential commonalities across primate species. Our data also provide evidence for a decrease in *FKBP5* DNA methylation across the normative development of these macaques between birth and 2 years of life. Across the observation period we detect a mean *FKBP5* DNA methylation difference of close to 25% (MALT: 41.12% ± 14.90 vs. CTRL: 65.75% ± 19.08) with marked differences between day 2 and month 6 and a convergence of DNA methylation levels at month 24 (**Figure 2**). This is in line with a developmental increase of cortisol secretion observed in humans and NHPs ^66,67^.

**Figure 2.**
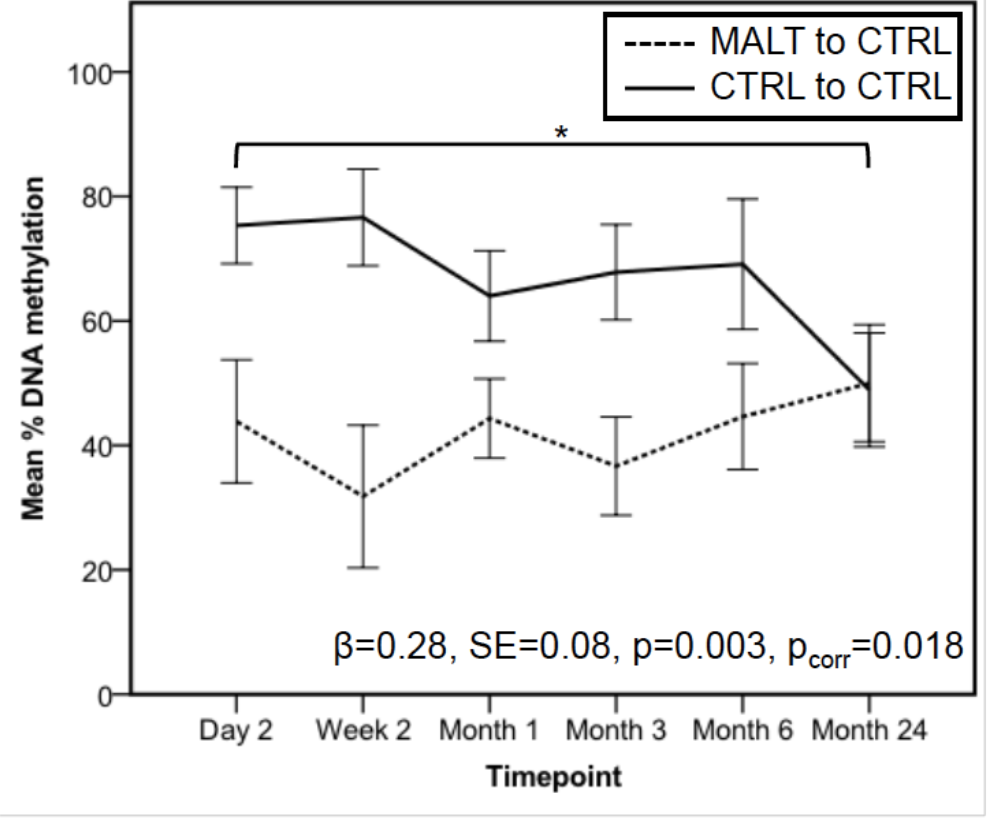
Group-dependent *FKBP5* DNA methylation differences across development. DNA methylation at previously described CpGs in and around functional glucocorticoid response elements (GREs) was determined using the Sequenom EpiTYPER platform across all 6 time points. MALT offspring (N=9) compared to CTRL offspring (N=12) show lower DNA methylation at CpG5 (β=0.28, SE=0.08, p=0.003, p_Bonferroni_=0.018). DNA methylation differences are pronounced between birth and month 6 and show a convergence at 2 years of age. Error bars indicate SEM.

### Functional impact of differential methylation within FKBP5 GRE binding sequence

We previously provided evidence for a functional role of DNA demethylation in and around GR binding sequences in human *FKBP5* intron 7 ^42^ with demethylation leading to a stronger GR binding to the intron 7 enhancer element, stronger transcriptional activation of *FKBP5*, thus underlying a dysregulation of the negative feedback on GR function and higher cortisol levels over time. Using a competitive ELISA assay, we now tested how DNA methylation at CpG5, which is located in the center of the GRE, affects binding of the GR in NHPs. Using 35mer synthetic oligonucleotides with either a methylated or unmethylated CpG5 cytosine, we found that the methylated GRE is less efficient in binding the GR compared to the unmethylated oligonucleotide in the macaque model (Mean absorbance 450/655nm, unmethylated=0.356 ± 0.035, methylated=0.476 ± 0.046, t(4)=−3.59, p=0.023). These data suggest that the differential *FKBP5* DNA methylation even at a single CpG site observed *in vivo* in rhesus monkeys could be associated with differential GR binding and GR function at the cellular level.

### DNA methylation of FKBP5 in biological mothers

We next tested the possibility of differential *FKBP5* DNA methylation in the adult biological MALT mothers in comparison to the biological CTRL mothers. However, we did not observe a significant difference in DNA methylation between MALT and CTRL biological mothers (Mean % DNA methylation ± SD, MALT: 90.38 ± 8.29, CTRL: 91.56 ± 9.30, t(15)=0.28, p=0.78) (**Supplemental Figure 2**).

### FKBP5 methylation and neuroendocrine outcomes in the offspring

Prior evidence suggests an important role of *FKBP5* methylation in the regulation of the HPA axis function, in particular in stress-reactive glucocorticoid negative feedback and active stress response ^34^. Using hair cortisol measurements at postnatal day 2 and 6 months of age, we aimed to assess *in utero* stress exposure of the newborns ^68^, and HPA activity during the first 6 postnatal months ^69^, respectively and relate our findings of differential *FKBP5* DNA methylation in the offspring to neuroendocrine regulation.

The lack of group differences on day 2 hair cortisol levels (log transformed pg/mg hair) suggest that a potential *in utero* exposure to higher stress levels in MALT newborns is less conceivable and does not explain lower *FKBP5* methylation levels in MALT newborns (MALT: 6.36 ± 0.28, CTRL: 6.28 ± 0.2, t(18)=−0.73, p=0.47) (**Supplemental Figure 3A**). No significant correlations were detected, either, between postnatal day 2 *FKBP5* DNA methylation and hair cortisol levels in either the entire sample nor the individual groups, which supports the hypothesis that prenatal *in utero* stress exposure is not the primary cause of reduced *FKBP5* DNA methylation in MALT newborns (MALT: r(7)=−0.18, p=0.70, CTRL: r(5)=0.35, p=0.56, Fisher z-score=−0.63, p=0.53).

We next tested the hypothesis that the observed differences in *FKBP5* DNA methylation exert an effect on postnatal HPA axis activity as measured by cortisol accumulation in hair from birth through month 6. Similar to our observations at day 2, there were no obvious group differences in hair cortisol levels between offspring from MALT and CTRL mothers (MALT: 4.74 ± 0.22, CTRL: 4.79 ± 0.24, t(16)=0.50, p=0.63) (**Supplemental Figure 3B**), which was expected given the non-MALT control environment provided by the CTRL foster mothers. However, *FKBP5* DNA methylation at month 6 showed an inverse correlation with month 6 hair cortisol between both groups (MALT r(8)=−0.76, p=0.031, CTRL: r(8)=0.69, p=0.06, Fisher z-score=−2.9, p=0.003) (**Figure 4**). This suggests a functional relationship between *FKBP5* DNA methylation and cortisol levels over time such that lower *FKBP5* DNA methylation in MALT offspring are associated with higher postnatal hair cortisol levels.

**Figure 3.**
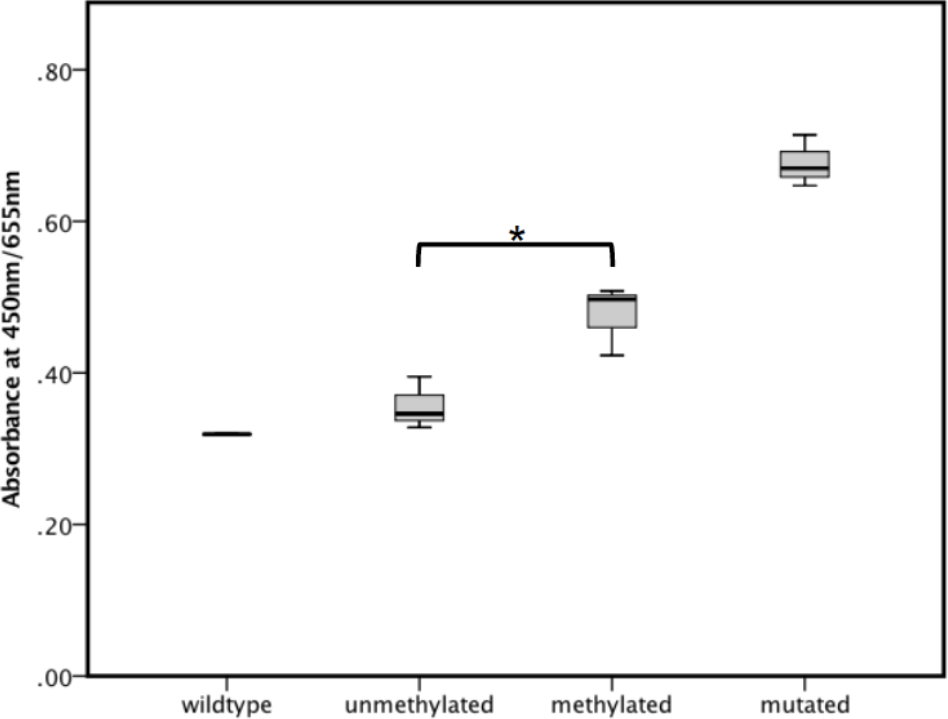
GR binding to methylated and unmethylated GRE sequence. To test the influence of DNA methylation within the GRE in intron 7 on GR binding, we performed a competitive binding enzyme-linked immunosorbent assay. A low absorbance indicates a strong binding of the competitive oligo to the GR and thus a reduction of available GR for detection via antibody binding and HRP-based color change. Vice versa, a high absorbance indicates a less efficient binding of the competitive oligo to the GR that hence binds to the consensus GRE sequences immobilized on the plate, leading to a strong calorimetric signal. The methylated CpG5 GRE is less efficient in binding the GR compared to the unmethylated oligonucleotide (Mean absorbance 450/655nm, unmethylated=0.356 ± 0.035, methylated=0.476 ± 0.046, t(4)=−3.59, p=0.023, N=3) indicating that a reduction of DNA methylation within the GRE lead to a stronger GR binding to this enhancer element.

**Figure 4.**
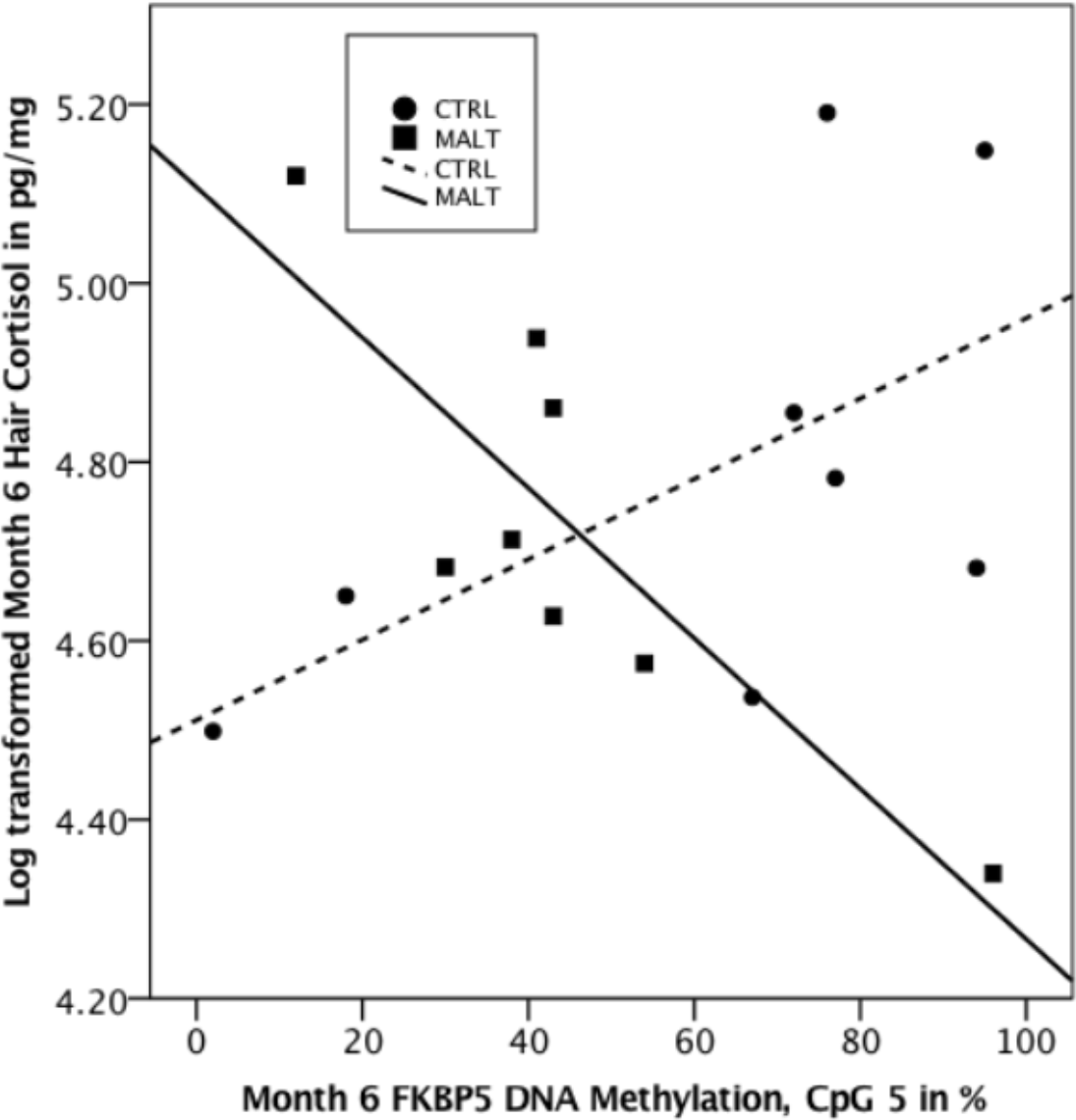
Correlation Month 6 DNAm and Month 6 Hair Cortisol in MALT and CTRL. A negative correlation between hair cortisol at 6 month and *FKBP5* DNA methylation was observed in MALT offspring (r(8)=−0.76, p=0.031), which is in line with previous reports providing evidence for a reduced *FKBP5* DNA methylation is leading to higher cortisol levels. In contrast, CTRL offspring showed a positive correlation between *FKBP5* DNA methylation and hair cortisol (r(8)=0.69, p=0.06). These data support the notion that *FKBP5* DNA methylation in MALT and CTRL offspring is influencing cortisol levels in opposite directions (Fisher z-score=−2.9, p=0.003) that may lead to functional differences in HPA axis activity under stressful conditions.

We corroborated these observations by analyzing associations with blood plasma cortisol levels. Neither day 2 baseline morning plasma cortisol (log transformed mean in ug/dl) (MALT: 1.91 ± 0.25, CTRL: 2.06 ± 0.36, t(15)=0.98, p=0.34) nor month 6 (MALT: 2.50 ± 0.17, CTRL: 2.37 ± 0.15, t(13)=−1.52, p=0.15) showed significant group differences. When calculating the mean plasma cortisol levels between day 2 and month 6 as a proxy for plasma cortisol levels over the first 6 months of life, we did not observe significant group differences either (MALT: 2.10 ± 0.37, CTRL: 2.18 ± 0.20, t(18)=0.65, p=0.53), which is congruent with our hair cortisol data. Correlations between day 2 plasma cortisol levels and day 2 *FKBP5* DNA methylation in MALT and CTRL offspring, were not statistically significant, either (MALT r(6)=−0.29, p=0.96, CTRL: r(4)=−0.2, p=0.8, Fisher z-score=−0.08, p=0.94). However, similar to our hair cortisol data, the cumulative plasma cortisol levels (day 2 to 6 months) significantly correlate with month 6 *FKBP5* DNA methylation in MALT offspring, and shows a significant difference between MALT and CTRL (MALT r(8)=−0.86, p=0.006, CTRL: r(10)=0.58, p=0.08, Fisher z-score=−3.34, p<0.001). These data further support the observation that MALT individuals with lower month 6 *FKBP5* DNA methylation have higher postnatal basal cortisol levels (i.e. HPA axis activation) over time.

In summary, cortisol data in hair and in plasma show no discernible group differences between offspring born to MALT vs. CTRL biological mothers, congruent with the competent maternal care both groups of infants experienced postnatally. However, offspring of biological MALT mothers that show reduced *FKBP5* DNA methylation levels show a significant negative correlation between DNA methylation and cortisol levels at month 6, but not at day 2 after birth, indicating that reduced DNA methylation may lead to increased cortisol levels postnatally. These data also suggest that although the baseline HPA axis activity is similar in MALT and CRTL offspring at birth, it may be differentially primed for responding to a stressful environment in the case of offspring of MALT biological mothers.

### FKBP5 methylation and behavioral outcomes in the offspring

Hair cortisol levels and basal morning plasma cortisol levels under non-stressful, resting conditions may not entirely reflect the influence of *FKBP5* DNA methylation on the dynamic regulation of the HPA axis. In absence of more direct readouts of the dynamic HPA axis regulation in our cohort, we aimed to investigate behavioral effects under mild stressful conditions with respect to ancestral phenotype and *FKBP5* DNA methylation. A broad body of literature shows robust effects of childhood maltreatment and ELS on a spectrum of behavioral outcomes in human and NHPs ^70–75^. We hypothesized that MALT offspring will show deficits in certain domains related to emotional and stress reactivity, compared to CTRLs. Therefore, we studied the infant’s behavioral reactivity at 6 months during the Human Intruder Test (HIT), a standard NHP paradigm that assesses emotional regulation in response to increasing environmental challenging conditions, and used to investigate stress and anxiety-related behavioral responses ^56,57,76^. Indeed, data from the HIT at 6 months suggests a functional effect of *FKBP5* methylation on behavioral outcomes, particularly on measures of anxiety, locomotion, freezing and vocalization during the ‘alone’, ‘profile’, and ‘stare’ conditions of the HIT ^57,76^. We found a significant group difference for locomotion during the ‘profile’ condition with MALT individuals showing increased locomotion compared to CTRL (log transformed mean % locomotion during 10 min observation period) (MALT: −2.42 ± 0.65, CTRL: −6.82 ± 3.86, t(17)=−3.7, p=0.004) (**Figure 5A**). Significant negative correlations were detected between *FKBP5* DNA methylation at month 6 and anxiety-like responses during the ‘alone’ condition (r(17)=−0.53, p=0.03), and anxiety behaviors and locomotion during the ‘profile’ condition (r(17)=−0.53, p=0.03 for anxiety and r(17)=−0.62, p=0.008 for locomotion), indicative of increased overall anxiety-like behaviors and locomotion in individuals with reduced *FKBP5* DNA methylation (**Table 2, Figure 5B**).

**Figure 5.**
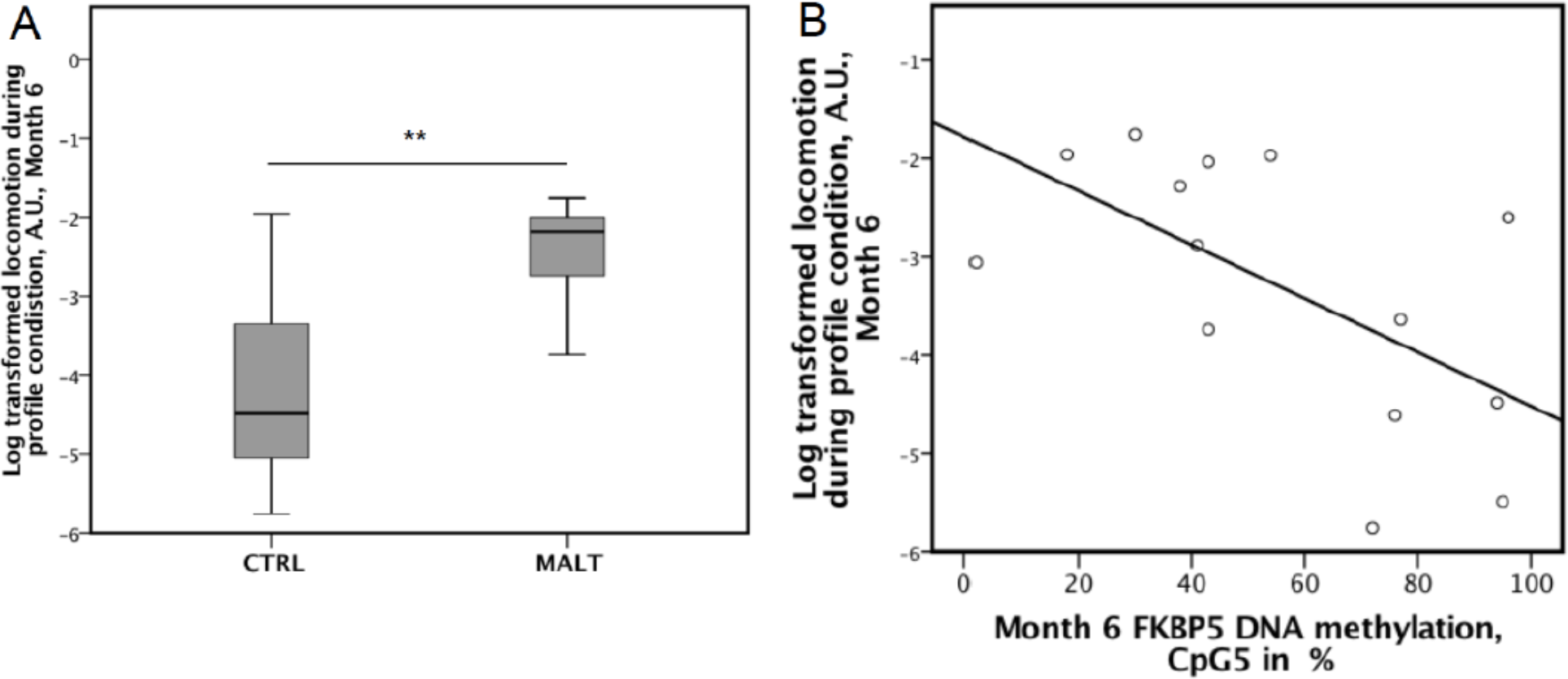
Locomotion during HIT profile condition between MALT and CTRL. **(A)** Boxplots showing a significant group difference for locomotion during the profile condition with MALT individuals showing increased locomotion compared to CTRL (log transformed mean % locomotion during 10 min observation period) (MALT: −2.42 ± 0.65, CTRL: −6.82 ± 3.86, t(17)=−3.7, p=0.004). **(B)** A negative correlation was detected between *FKBP5* DNA methylation at month 6 and anxiety during the alone condition (r(17)=−0.53, p=0.03), and anxiety and locomotion during the profile condition (r(17)=−0.53, p=0.03 for anxiety and r(17)=−0.62, p=0.008 for locomotion), which is indicative of increased overall anxiety and locomotion in individuals with reduced *FKBP5* DNA methylation under stressful conditions.

**Table 2.** Spearman’s rank correlation of *FKBP5* methylation and behavioral phenotypes during the Human Intruder Test in the offspring at month 6. Table showing the alone (left), profile (middle) and stare (right) conditions and correlations between month 6 *FKBP5* methylation and functional behavioral readouts. Negative correlations between DNA methylation and anxiety and locomotor activity indicates that reduces DNA methylation in our cohort is associated with increased levels of anxiety and locomotion.

|  |  | Alone |  |  |  | Profile |  |  |  | Stare |  |  |  |
| --- | --- | --- | --- | --- | --- | --- | --- | --- | --- | --- | --- | --- | --- |
|  |  | Anx<sup>(1)</sup> | Loc<sup>(2)</sup> | Frz<sup>(3)</sup> | Coo<sup>(4)</sup> | Anx<sup>(1)</sup> | Loc<sup>(2)</sup> | Frz<sup>(3)</sup> | Coo<sup>(4)</sup> | Anx<sup>(1)</sup> | Loc<sup>(2)</sup> | Frz<sup>(3)</sup> | Coo<sup>(4)</sup> |
| **Month 6**<br>***FKBP5***<br>**DNA methylation** | Spearman's rho | -.529 | -0.043 | 0.243 | -0.385 | -.532 | -.621 | 0.230 | -0.422 | -0.151 | 0.429 | 0.418 | -0.362 |
|  | Sig. 2-tailed | **0.029** | 0.870 | 0.348 | 0.127 | **0.028** | **0.008** | 0.375 | 0.091 | 0.562 | 0.086 | 0.095 | 0.154 |
|  | N | 17 | 17 | 17 | 17 | 17 | 17 | 17 | 17 | 17 | 17 | 17 | 17 |

Footnotes: <sup>(1)</sup> Anx=Anxiety; <sup>(2)</sup> Loc=Locomotion; <sup>(3)</sup> Frz=Freezing; <sup>(4)</sup> Coo=Cooing

## Discussion

The transmission of environmental information and experiences across generations and their importance for relevant molecular, physiological, behavioral and medical phenotypes remains a critical, yet still controversial question. Although recent studies in humans may suggest inter- and even transgenerational transmission or inheritance of acquired phenotypes, the limitations of current experimental designs and approaches prevent more definitive answers as to what extent the environment can influence phenotypes across generations and what the underlying mechanisms are in primates.

In this study, we leveraged a unique, well-controlled randomized cross-fostering design in rhesus monkeys, where newborn macaques from infant maltreating or competent, nurturing care lineages were adopted and raised by competent, nurturing foster mothers. Importantly, maltreatment appears to be a stable and natural phenotype discernible in captivity and in the wild, which is not provoked through experimental conditions and is thus well suited as a translational model for childhood maltreatment in humans. The cross-fostering design enables us to specifically investigate the influence of the ancestral experience of the biological mother on molecular, neuroendocrine, and behavioral outcomes in the offspring. In our studies, offspring were cross-fostered immediately at birth, and animals grew up under competent maternal care conditions, not exposed to maltreatment and without contact with the MALT biological mother, thus preventing behavioral transmission of information. Moreover, the longitudinal design of this study allowed us to investigate molecular phenotypes across developmental trajectories in the offspring from birth through the juvenile, prepubertal period. Of note, the design excluded major confounding factors as both groups were balanced with regard to sex of the offspring and age, rank, parity, and gravity status of both the biological and foster mothers. In addition, all individuals shared a similar environment with regard to housing, nutrition, veterinary care, and exposure to natural conditions at the Yerkes National Primate Research Center (YNPRC) Field Station in Lawrenceville, GA.

We show that ancestral maltreatment in the biological mother leads to reduced *FKBP5* DNA methylation in offspring at birth and during the infant period, providing evidence for an intergenerational effect of childhood maltreatment on epigenetic programming in this NHP primate model. These results are in line with our observations in humans exposed to sexual and physical maltreatment during childhood ^42^ suggesting comparable programming of DNA methylation in *FKBP5* in response to childhood maltreatment across species with similar spatial and directional effects. Moreover, studies in rodents exposed to glucocorticoids demonstrated similar results suggesting conserved mechanisms across a wide range of species ^77^. Infant macaques born to biological MALT mothers showed lower *FKBP5* DNA methylation levels than individuals born to biological CTRL mothers, a difference that is already present at day 2 after birth and lasting at least until 6 months of age (**Figure 2**). The convergence of MALT and CTRL offspring methylation profiles may reflect a compensatory role of the control postnatal environment in which all offspring monkeys were raised. The CpGs found demethylated in prior human studies ^26,35,42,78–80^ and in this report are spatially central to a functional enhancer element that harbors glucocorticoid response elements (GREs) in intron 7 of *FKBP5*. These GREs regulate the dynamic transcriptional response of *FKBP5* and thus HPA axis activity, the stress response and glucocorticoid negative feedback ^26,34,35,42,78,79,81,82^. Our findings in macaques are consistent with that literature, showing that DNA methylation in the central CpG5 position of the *FKBP5* intron 7 GRE in the macaque gene influence GR binding *in vitro*, and are subsequently associated with differential FKBP5 regulation of GR feedback and peripheral cortisol levels.

Although we cannot definitively exclude the influence of underlying genetic variants in these monkeys, several aspects make this assumption less likely. First, the behavioral phenotype of infant maltreatment in these monkeys is genotype independent, cross fostering effectively stops the transmission of this behavior, excluding a strong genetic effect. Second, individuals included in this study came from different genetic backgrounds as part of the overall breeding scheme of the YNPRC. Third, for a fixed genetic effect, we would assume that DNA methylation differences are rather stable over time and across generations, which is not the case in our data. When testing the biological mothers for their *FKBP5* DNA methylation profiles, we did not observe significant differences. This indicates that the differences we detect during the first months of development in the offspring may not be stable across life, which is in line with our observation that methylation differences between groups show no difference at 24 months. This observation is in contrast to findings in humans where differences in *FKBP5* DNA methylation appear to remain stable into adulthood ^35,42^. However, human studies typically involve an environment that remains similar throughout life. Individuals exposed to childhood maltreatment often continue to live in stressful environments as reflected by the subsequent exposure to adult trauma and other factors such as low socioeconomic status ^83,84^. Data presented here in fact support the notion that a non-stressful postnatal environment (in this case, competent maternal care postnatally) may mitigate the intergenerational transmission of risk in form of *FKBP5* DNA methylation as seen by the end of the study period where DNA methylation differences disappear between offspring of biological MALT and CTRL mothers. This is also in line with previous observations suggesting that cross-fostering offspring from MALT biological mothers to CTRL foster mothers inhibit the transmission of maltreatment across generations ^59^. This is indicative of a non-genetic intergenerational transmission of this particular behavior. Our data show that there are, however, other long-term molecular, neuroendocrine and behavioral alterations detectable in offspring born to MALT in contrast to CTRL lineages, independent of postnatal maternal care received. Nevertheless, these differences seem not sufficient to induce the MALT phenotype.

DNA methylation in *FKBP5* has been associated with the function of the HPA axis, dynamic cortisol release and negative feedback signaling of *FKBP5* towards the glucocorticoid receptor. We previously provided evidence for a negative correlation of *FKBP5* DNA methylation and cortisol levels in line with a dysregulation of the HPA axis in humans exposed to trauma or severe stress ^42^. Thus, we hypothesized that regulation of the HPA axis is altered in MALT offspring compared to CTRL. Hair cortisol concentrations are indicative of prolonged cortisol accumulation and provide a retrospective reflection of integrated cortisol secretion over periods of several months in humans and non-human primates ^69,85^. In addition, hair cortisol levels in newborn monkeys are reflective of *in utero* stress exposure ^68^. We used hair cortisol collected on day 2 after birth to test whether prenatal stress could explain the lower *FKBP5* methylation in offspring from MALT lineages compared to CTRL offspring. However, the lack of differences between groups ruled out this possibility. This may also point to a mechanism of transmission that includes early effects of maltreatment on the germline cell biology; however, we cannot exclude the possibility of *in utero* stress effects independent of our hair cortisol readout. Lower *FKBP5* DNA methylation levels in humans are associated with less effective negative feedback on the HPA axis and higher cortisol levels over time ^26,42,86,87^. In line with these observations, a negative correlation of *FKBP5* DNA methylation at month 6 with elevated HPA axis activity as measured by both elevated hair and plasma cortisol levels at 6 months was observed in offspring from MALT biological mothers. These findings support prior evidence of disturbed feedback mechanisms of *FKBP5* on the GR and provide evidence for important functional effects of *FKBP5* DNA methylation differences in the rhesus offspring. It is important to note that we did not detect group differences in hair and plasma cortisol at month 6, which is actually in line with the function of *FKBP5* as a regulator of the *dynamic* feedback after stress exposure. Since MALT and CTRL offspring are raised in control environmental conditions, we would not expect to be able to detect baseline differences due to a lack of major postnatal stressful conditions for both groups. However, to test the dynamic response of MALT and CTRL offspring to a stressful event, we choose the Human Intruder Test (HIT) a mildly stressful task, similar to the Strange Situation Test used in human children ^76,88^ that assesses the response to changing environmental conditions and induces stress and anxiety-related behaviors. Testing the NHP offspring in the ‘alone’ condition primarily assesses anxiety-related and vocalization phenotypes whereas the ‘profile’ and ‘stare’ conditions evaluate aggressive behaviors, locomotion and freezing. In line with prior evidence suggesting an exaggerated response to the HIT in monkeys with higher cortisol levels ^89^, we show a negative correlation between *FKBP5* methylation at month 6 and anxiety-like behavior during the ‘alone’ condition as well as anxiety and locomotion during the ‘profile’ condition. This suggests increased overall anxiety and locomotion in individuals with reduced *FKBP5* DNA methylation, which is in agreement with the hypotheses that lower *FKBP5* methylation, would lead to higher cortisol levels during a stressful task.

Several human studies support a role for *FKBP5* DNA demethylation in response to ELS ^35,42,78–80^. Importantly, a study in survivors of the Holocaust suggested additional intergenerational effects of trauma exposure on *FKBP5* DNA methylation in the offspring ^26^. The Holocaust-exposed parental generation showed increased DNA methylation whereas the offspring generation showed decreased *FKBP5* DNA methylation levels in and around the orthologous intron 7 GREs ^26^. In addition, lower levels of DNA methylation were negatively correlated with cortisol awakening response, which in general is in line with our observations. As mentioned before, human studies investigating such phenomena are incredibly difficult to control and not surprisingly provoke strong excitement and controversy at the same time. Similar to human studies, as yet we are unable to provide mechanistic insight to the phenomena we report in this study with NHPs at the moment. However, we emphasize that the infant maltreatment model in rhesus monkeys used in this study is highly translational, and well-controlled, thus providing cross-species supportive evidence for intergenerational effects of maltreatment in childhood, one of the most critical risk factors for psychiatric disorders. This model is optimally suited for further understanding of mechanisms of intergenerational transmission of information in primates.

Overall, our data suggest a role for intergenerational programming of epigenetic patterns in response to childhood maltreatment in primates, independent of behavioral transmission. The observed differences in *FKBP5* methylation show functional effects on HPA axis activity and behavioral outcomes. Thus, childhood maltreatment in primates may induce molecular and behavioral changes in a subsequent generation leading to a higher risk for disease across generations.

## Methods

### Subjects and Experimental Groups

We used a unique and innovative cross-fostering design with random assignment to experimental group at birth. To investigate the contribution of ancestral maltreatment history on the next generation *FKBP5* methylation and related neuroendocrine and behavioral phenotypes, we focused on infants born to maltreated (MALT) or control (CTRL) biological mothers that were randomly assigned and cross-fostered at birth to control foster mothers with a history of nurturing maternal care (CTRL) following previous protocols ^58^, thus creating two separate groups: CTRL-to-CTRL and MALT-to-CTRL (**Figure 1**). Thus, we are able to ask specifically how maltreatment in the biological mothers influences offspring outcomes eliminating the possibility of behavioral effects from the biological mother towards the offspring, including variations in maternal care, as offspring grew up in a control environment different from the biological mother. A total of N=20 infant rhesus monkeys (*Macaca mulatta*), 9 males and 11 females, and their biological and foster mothers, served as subjects for the proposed studies, with 11 infants assigned at birth to the CTRL-to-CTRL group (6 males, 5 females), and the remaining 9 infants assigned to the MALT-to-CTRL group (3 males, 6 females). (**Table 1, Figure 1**). Biological and behavioral data were collected longitudinally in the offspring from birth to 24 months of age (juvenile, prepubertal period). All subjects lived with their foster mothers and families in large, complex, social groups maintained at Emory University’s Yerkes National Primate Research Center (YNPRC) Field Station in Lawrenceville, GA. These social groups consisted of 75-150 adult females, their sub-adult and juvenile offspring and 2-3 adult males, and allowed us to counterbalance our experimental groups for sex, social dominance rank (high, medium and low social status), maternal parity/gravity/age, and to select subjects from different matrilines (i.e. to be genetically unrelated). Animals were housed in 100ftx100ft compounds with access to climate controlled indoor areas. Standard nutrition (Purina Mills Int., Lab Diets, St. Louis, MO) and seasonal fruits and vegetables were provided twice daily with water access *ad libitum*. All procedures were in accordance with the Animal Welfare Act and the US Department of Health and Human Services “Guide for the Care of Laboratory Animals” and approved by the Emory University Institutional Animal Care and Use Committee (IACUC).

### Behavioral characterization of maternal care

Maternal care is a stable and consistent trait across generations with abusive behavior transmitted from mothers to daughters, resulting in maternal lineages showing infant maltreatment or competent maternal care, respectively ^45,90^. We study these lineages in our colony for several years. We selected the CTRL and MALT biological mothers based on their consistent maternal behavior towards previous offspring as described in detail below. In order to characterize their own exposure to maltreatment, we tracked direct experimental observations by our lab and colony records to confirm that the biological MALT mothers were maltreated themselves as infants. This was available for 6 out of 9 biological MALT mothers. Based on the consistency of maltreatment across generations, the remaining 3 biological MALT mothers were defined as maltreated based on the consistent maltreating phenotypes of their mothers. The absence of any records of maltreatment towards the CTRL biological mothers defined the CTRL biological mothers who themselves did not have any records of maltreatment towards previous offspring. All foster mothers in this study were selected based on their consistent competent maternal care towards prior offspring, which was confirmed through direct observations of their maternal behavior towards fostered infants. A detailed description of the behavioral characterization of competent maternal care (CTRL) in contrast to infant MALT is provided in previous publications ^45,48,53,55,58,91^. Briefly, focal observations of maternal care are performed over the first 3 months of the infants life (30 min long observation on separate days: 5 days/week during month 1, 2 days per week on month 2 and 1 day/week during month 3, for a total of 16 hours. This observation protocol is optimal to document early maternal care in this species, particularly given that physical abuse is the highest during month. Competent maternal care (CTRL) was defined as typical behaviors such as nursing, cradling, grooming ventral contact and protection of the infant. Maltreatment (MALT) was defined as the comorbid occurrence of physical abuse (operationalized as violent behaviors directed towards the infant that cause pain and distress, including dragging, crushing, throwing) and infant rejection (i.e. prevention of nipple/ventral contact and pushing the infant away) resulting in high levels of infant distress ^45,52–55^.

### DNA collection

Genomic DNA for *FKBP5* DNA methylation analysis was obtained from whole blood consistently collected during the morning hours in EDTA tubes (Sarstedt, Newton, NC) at the following time points during development: Day 2, Day 14 (Week 2), Day 30 (Month 1), Day 90 (Month 3), Day 180 (Month 6), Day 720 (Month 24). DNA was extracted using standard procedures. We also obtained DNA samples from all biological mothers (N=20).

### *FKBP5* DNA methylation analysis

Based on our previous work on human *FKBP5*, we adapted the rhesus *FKBP5* DNA methylation assay from our human assays interrogating DNA methylation in and around functional glucocorticoid response elements (GREs) in intron 7 of *FKBP5* ^42^. We interrogated 6 CpG dinucleotides that are orthologous to the human CpGs investigated before. Briefly, at Varionostic GmbH (Ulm, Germany) genomic DNA was bisulfite converted using the EZ-96 DNA Methylation Kit (Zymo, Irvine, CA). Amplicons, 476bp in length were generated from bisulfite converted DNA using primer P1_F 5’-aggaagagagGTTGTTTTTGGAATTT AAGGTAATTG-3’ and P1_R 5’-cagtaatacgactcactatagggagaaggctTCTCTTACCTCCA ACACTACTACTAAAA-3’. PCR reactions contained 10x buffer, HotStarTaq, MM, containing 1,5 mmol Mg2+, dNTPs (Qiagen, Hilden, Germany) (5ul), 5 μM Forward primer (0.5ul) 5 μM Reverse primer (0.5ul), Bisulfite DNA (1ul), H2O (3ul). PCR cycling conditions were [95C-15’00’’, 49x (95C-40’’, 58C-40’’, 72C-40’’), 72C-5’00’’, 4C]. Amplicons were then analyzed on the Sequenom EpiTYPER platform in accordance with the manufacturers recommendations and included control standards and non-template control (please see **Supplemental Figure 1** for rhesus *FKBP5* intron 7 assay in comparison to the human intron 7 assay).

### Enzyme-linked immunosorbent assay (ELISA)

To test the influence of DNA methylation within the GR binding site in intron 7 on the binding capabilities of GR to the putative GRE, we performed a competitive binding enzyme-linked immunosorbent assay using the TransAM GR kit (catalog no. 45496; Active Motif, Carlsbad, CA). To each microwell coated with canonical GRE sequence oligonucleotides, we added dexamethasone-treated HeLa nuclear extract (5 μg; Active Motif) and 40 pmol of one of four double-stranded oligo DNAs (two competing/experimental sequences plus a negative and positive control). All assays were performed in triplicate. Individual oligos and nuclear extract were added to each microwell and incubated for 1 h. Next, the primary antibody (GR 1:1000) and then secondary antibody (horseradish peroxidase (HRP) - conjugated IgG; 1:1000) was added for 1 h each. After several rinses, a colorimetric reaction to horseradish peroxidase was initiated and stopped after 7minutes. Absorbance was measured at a wavelength of 450 nm and 655 nm to determine binding efficiencies. Low absorbance indicates a strong binding of the competitive oligo to the GR and thus a reduction of available GR for detection via antibody binding and HRP-based color change. Vice versa, a high absorbance indicates a less efficient binding of the competitive oligo to the GR that hence binds to the consensus GRE sequences immobilized on the plate, leading to a strong calorimetric signal. Antibodies, positive/negative controls and other reagents were provided in the TransAM GR kit. Custom methylated primer sequences: mGRE-fwd 5’-TCCTGTGAAGGGTACAATC/iMe-dC/GTTCAGCTCTGAAAA-3’ and mGRE-rev 5’-TTTTCAGAGCTGAA/iMe-dC/GGATTGTACCCTTCACAGGA-3’. Custom unmethylated primer sequences: GRE-fwd 5’-TCCTGTGAAGGGTACAATCCGTTCAGCTCTGAAAA-3’ and GRE-rev 5’-TTTTCAGAGCTGAACGGATTGTACCCTTCACAGGA-3’ (all IDT Technologies).

### Behavioral Data

Emotional reactivity of the offspring to varying degrees of threat was examined at 6 months of age during the Human Intruder Test (HIT; ^76^). On the testing day, the trained mother-infant pairs were accessed in their home compound and the infant transported to a novel testing room and placed alone in a testing cage with one side made of clear plexiglass to allow video recording following published procedures ^57,58^. The HIT is widely used to assess emotional reactivity (including fearful, anxious and aggressive behaviors) to an unfamiliar human in rhesus monkeys under three different and consecutive conditions, 10 min each, that pose an increasing degree of threat: 1) ‘Alone’ condition (no human intruder), which elicits “calling” vocalizations (coos), locomotion and exploration aimed to escape; 2) next, an unfamiliar human intruder entered the room and stood 3m from the cage presenting his/her profile to the subject without direct eye contact (‘profile’ condition), which poses an uncertain threat that results in behavioral inhibition (freezing behavior), stop of vocalizations and vigilance; and 3) direct eye contact of the intruder with the monkey (‘stare’ condition), which is a threatening behavior for these animals, which typically stop freezing and respond with aggressive and submissive behaviors and threat vocalization towards the intruder. Immediately following the 30 min HIT, the infants were reunited with their mothers and returned to their social groups. All sessions were video recorded and later scored by two trained coders (inter-rater reliability of >90% agreement) using The Observer XT software (v10.5, Noldus, Inc., Netherlands) and published ethograms and procedures ^57,58^. For this study, we focused on the analysis of anxiety, freezing, cooing and locomotion behaviors in line with the role of *FKBP5* and the HPA axis in these behaviors.

### Hair Cortisol

Hair cortisol data were obtained as described before ^55,85^. Briefly, approximately one square inch area of hair was shaved from the nuchal area of the head on postnatal day 2 after birth (to measure cortisol accumulated in hair *prenatally*) and at month 6 (to measure cortisol accumulated in hair that grew between birth and 6 months). The hair was weighed, washed with isopropanol, ground to fine powder and extracted with methanol overnight. After methanol evaporation, the residue was re-dissolved in assay buffer and cortisol was determined using the cortisol ELISA kit (Salimetrics, #1-3002; Carlsbad, CA). Intra- and inter-assay coefficients of variation were <10%.

### Morning Plasma Cortisol

Procedures for collection of morning basal blood samples have been described before ^67,90^. Blood samples were collected at the same ages as hair cortisol (postnatal day 2, and 6 months) in EDTA tubes and immediately placed on ice. Plasma was separated by centrifugation at 1,000g for 10 min at 4°C and then aliquoted and stored at −80°C until assayed. Plasma concentrations of cortisol were assayed by ultra-performance liquid chromatography electrospray ionization (UPLC-ESI) tandem mass spectrometry in duplicates by the YNPRC Biomarkers Core, following published methods^92^.

### Data Analyses

Demographic and descriptive data on the sample are given in **Table 1** separated for biological MALT and CTRL groups. Groups show a balanced design with respect to major potential confounding variables (e.g. maternal age, parity, social rank, offspring sex). Between group differences for demographic variables in **Table 1** were determined using independent samples t-tests or chi-square tests. A simple linear mixed effects model was used to investigate differences between ancestral groups (MALT and CTRL), in terms of the offspring overall mean *FKBP5* DNA methylation levels across all time points (day 2, week 2, month 1, month 3, month 6 and month 24); this comparison of means is very similar to a comparison of the area under the curve (AUC approach). *FKBP5* DNA methylation was set as the dependent variable with 6 repeated measures over time; group (MALT, CTRL) was included as a fixed effect and was the effect of primary interest. To account for correlation among the 6 repeated measures and possibly heterogeneous variability over time, the model incorporated a random effect for individual (subject) and separate error variances at the 6 time points. Given our balanced study design with regard to potential confounding factors (biological and foster mother age, rank, parity, gravity; offspring sex), we did not include these variables in the final model, although we first ran preliminary analyses where each of them were separately tested in the model to ensure that they had, indeed, no major influence on the results of the analysis. Linear correlations between variables were performed using Spearman’s rank correlation or Pearson’s product-moment correlation (two-sided). All analyses were performed in SPSS 24. We used the Shapiro-Wilk Test to confirm normality and log transformed data that deviated from normal distribution. Significance was determined as p<0.05. Correction for multiple testing according to Bonferroni was applied for the number of CpGs tested.

## Acknowledgments

This work was supported by the following sources of funding: the National Institute of Mental Health (P50 MH078105 and MH086203 to BRH), the National Institute on Drug Abuse (R01 DA038588-01 to MMS), the Eunice Kennedy Shriver National Institute of Child Health and Human Development (R21 HD088931-01A1 to MMS, KR and TK) the Brain & Behavior Research Foundation (NARSAD YI 20895 to TK), EMBO (EMBO ALTF 1153-2013 to TK) and Office of Research Infrastructure Programs/OD Grant OD11132 (Yerkes National Primate Research Center –YNPRC-Base Grant, formerly RR000165). The authors thank Anne Glenn, Christine Marsteller, Zach Johnson and the staff at the YNPRC for the excellent technical support and animal care provided during the studies. The YNPRC is fully accredited by AAALAC, International.

## Author contributions

TK, KJR and MMS designed research; TK, EM, BRH, SB, DG, JSM and MMS performed research; TK, EM, BRH, DG, KJR and MMS analyzed data; and TK and MMS wrote the paper. All authors commented and approved the paper.

## Supplementary Materials

Supplementary Figures: S1-S3

